# EEG correlates of active removal from working memory

**DOI:** 10.1101/2025.01.09.632207

**Authors:** Jiangang Shan, Bradley R. Postle

## Abstract

The removal of no-longer-relevant information from visual working memory (WM) is important for the functioning of WM, given its severe capacity limitation. Previously, with an “ABC-retrocuing” WM task, we have shown that removing information can be accomplished in different ways: by simply withdrawing attention from the newly irrelevant memory item (IMI; i.e., via “passive removal”); or by or “actively” removing the IMI from WM (Shan and Postle, 2022). Here, to investigate the neural mechanisms behind active removal, we recorded electroencephalogram (EEG) signals from human subjects (both sexes) performing the ABC-retrocuing task. Specifically, we tested the hijacked adaptation model, which posits that active removal is accomplished by a top-down-triggered down-modulation of the gain of perceptual circuits, such that sensory channels tuned to the to-be-removed information become less sensitive. Behaviorally, analyses revealed that, relative to passive removal, active removal produced a decline in the familiarity landscape centered on the IMI. Neurally, we focused on two epochs of the task, corresponding to the triggering, and to the consequence, of active removal. With regard to triggering, we observed a stronger anterior-to-posterior traveling wave for active versus passive removal. With regard to the consequence(s) of removal, the response to a task-irrelevant “ping” was reduced for active removal, as assessed with ERP and with posterior-to-anterior traveling waves, suggesting that active removal led to decreased excitability in perceptual circuits centered on the IMI.

**Significance Statement:** The removal of no-longer-relevant information from working memory is critical for the flexible control of behavior. However, to our knowledge, the only explicit accounts of this operation describe the simple withdrawal of attention from that information (i.e., “passive removal”). Here, with measurements of behavior and electroencephalography (EEG), we provide evidence for a specific mechanism for the active removal of information from WM–hijacked adaptation–via the top-down triggering of an adaptation-like down-regulation of gain of the perceptual circuits tuned to the to-be-removed information. These results may have implications for disorders of mental health, including rumination, intrusion of negative thoughts, and hallucination.

## Introduction

A hallmark property of working memory (WM) is that it is rapidly updatable, in that new information can be added “on the fly” as required by moment-to-moment changes in the environment and/or in behavioral goals. And because of the severe capacity limitation of WM, this updating process is commonly assumed to also entail the removal of no-longer-relevant information. Despite this, however, empirical evidence indicates that removal is often not absolute. For example, the recall (e.g., of the orientation of a grating) of the current trial’s item is often biased toward the sample from the previous trial (i.e., an attractive serial bias, e.g., (Fischer and Whitney, 2014; Samaha et al., 2019)). On the neural level, there is also considerable evidence for the incomplete removal of the no-longer-relevant information, with information from the previous trial persisting into the current trial (Bae and Luck, 2019; Barbosa et al., 2020; Zhang and Lewis-Peacock, 2024). All these findings suggest that, under many circumstances, updating may involve a default strategy of “passive removal” (i.e., the simple withdrawal of attention from no-longer-relevant information), such that some residual trace of this information remains in the cognitive system, and can interfere with subsequent behaviors.

There is, however, evidence that updating can involve the active removal of information from WM. In a behavioral study using an “ABC-retrocuing” WM task, whereas passive removal resulted in an attractive serial bias (replicating, e.g., (Fischer and Whitney, 2014; Bliss et al., 2017; Samaha et al., 2019) active removal had the opposite effect: a repulsive serial bias. Similarly, in a dual WM+discrimination task, the condition encouraging passive removal produced attractive serial dependence, but the condition encouraging active removal produced repulsive serial dependence (Teng et al., 2022). These reversals of the sign of the serial dependence effect suggest that active removal may entail the transformation of the IMI into a “reversed” version of itself, one that also reverses its influence on subsequent WM processing.

This idea gains support from a functional magnetic resonance imaging (fMRI) study of WM with a retrocuing (Lorenc et al., 2020), the multivariate representation of a no-longer-relevant item was reversed after it acquired this status. Computational modeling indicated that this result was best explained by a mechanism in which the gain of perceptual circuits tuned to this item was down-modulated (Lorenc et al., 2020), a mechanism reminiscent of the one underlying sensory adaptation (Jin et al., 2005).

The repulsive bias of serial dependence is believed to arise during the encoding of the current trial’s item, due to sensory adaptation (Fritsche et al., 2017; Pascucci et al., 2019; Fritsche et al., 2020; Sheehan and Serences, 2022). Together, these findings have led us to speculate that the repulsive effects associated with active removal may implicate an adaptation-like mechanism. We refer to this mechanism as the “hijacked adaption” model of active removal from WM.

The current study was carried out to test behavioral and neural predictions of the “hijacked adaptation” model, whereby the active removal of information in WM is accomplished by a top-down modulation of the perceptual channels tuned to that information. To do this we recorded EEG activity from subjects performing an ABC-retrocuing WM task and focused on two epochs in the trial to assess the triggering of the active removal process, and, later in the trial, the consequence(s) of its deployment.

## Materials and Methods

### Subjects

Twenty-seven subjects from the University of Wisconsin–Madison community participated in the study and were compensated monetarily. All subjects provided informed consent approved by the University of Wisconsin–Madison Health Sciences Institutional Review Board. One was removed from analyses for failing to follow task instructions; another was removed due to poor performance (mean absolute recall error more than 2 SD higher than the group average). Thus, data from 25 subjects (19 females, 6 males, mean age = 23.8, range = 19 - 30) were included in all analyses.

### Stimuli

Subjects were seated in a dimly lit room at a viewing distance of 50 cm from the monitor. A chin rest was used to help keep the head stable during the task. Memory-sample stimuli were oriented gratings (radius = 3°, spatial frequency = 1circle/°, contrast = 0.5, random phase) presented at six possible locations (30°, 90°, 150°, 210°, 270°, 330°) on an imaginary circle with radius of 7°. Stimuli were white (RGB = 255,255,255) appearing on a gray (128,128,128) background. Ping stimuli were white bullseyes presented at all six locations (radius = 3°, spatial frequency = 1circle/°, contrast = 1, random phase).The ping was intended to provide strong but orientation-neutral stimulation of circuits involved in visual perception. Retrocues were white circles with the same radius as the samples, and the probes were black response dials (unfilled black circles with a black line corresponding to the diameter of the circle) with the same radius and random starting orientation. A white fixation dot was presented at the center of the screen throughout each block of trials.

### Experimental design

Subjects completed 6 blocks of ABC-retrocuing task (Fig. 1) in one session, 3 blocks of “no-overlap” trials followed by 3 blocks of “overlap” trials. Each block had 120 trials and lasted for ∼27 min. Subjects were asked to take a self-paced pause every 40 trials (i.e., ∼ 9 min) and were required to keep their head stable during the pause. Each trial started with central fixation (750 ms), after which two sample gratings (items *A* and *B*) were presented at different locations (1000 ms). Subjects were to remember these two samples across an initial delay (Delay 1; 1500 ms), after which a retrocue appeared at the location occupied by one of the two gratings (750 ms), thereby indicating that this item might be tested at the end of the trial. (Subjects were explicitly informed that the uncued item would not be tested, thereby making it an IMI.).After a second delay (Delay 2.1; 2000 ms), the ping was presented for 250 ms, followed by Delay 2.2 (1000 ms), then another sample grating (item *C*, 500 ms), Delay 3 (1000-ms), and finally a probe that appeared at the location that had been occupied by the retrocued item or by item C (with equal chance). The probe was displayed for 3000 sec, during which the subject was to recall the orientation of the probed item by adjusting the orientation of the response dial with a computer mouse, followed by feedback displaying the number of degrees of error (1000 ms). The inter-trial interval varied randomly from 500 to 700 ms.

**Figure 1.**
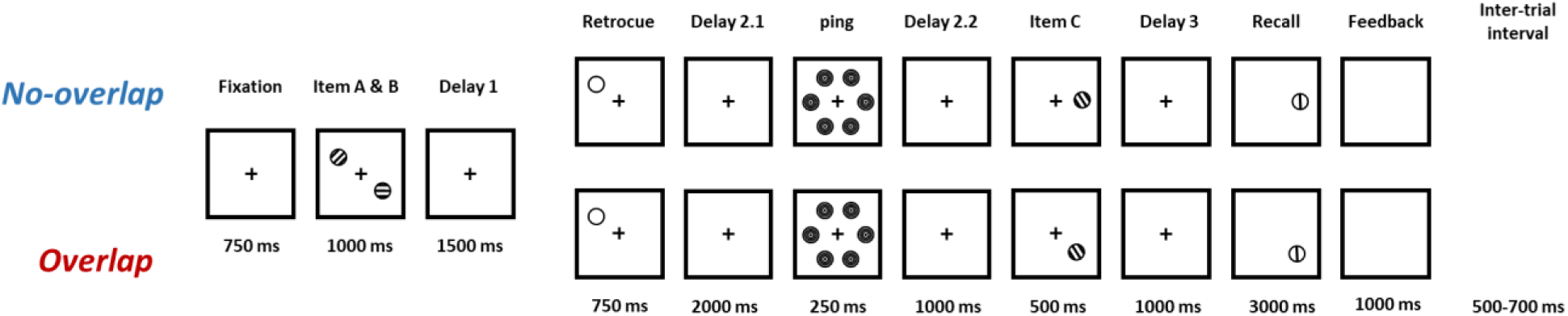
The experimental procedure.

The orientations of items *A*, *B*, and *C* were randomly drawn, with replacement, from a pool of 6 base orientations (20°, 50°, 80°, 110°, 140°, 170°), with a random jitter of -3 to 3° added. The locations of *A* and *B* were randomly selected from two of the six possible locations. Importantly, for the first three blocks (the no-overlap condition), item *C* was randomly presented at one of the four locations that had not been occupied by *A* or *B*, a fact that was specified during pre-experiment instructions. For the final three blocks (the overlap condition item), *C* was always presented at the same location as had been the IMI. The logic of this manipulation was that, because the overlap condition featured higher cue conflict between the IMI and item *C*, subjects would be motivated to actively remove the IMI from WM as soon as it was so designated by the retrocue. For the no-overlap condition, in contrast, because item *C* was always presented at a new location, the potential interference from IMI would be weaker, and so subjects might just use the default strategy of passive removal of IMI. Note that there was no explicit instruction about using active or passive removal. The location-related difference between the overlap and the no-overlap condition was only explained to subjects after they had completed the first three blocks of the experiment, a procedural detail intended to reduce the likelihood that subjects would engage an active removal strategy on no-overlap trials.

### Behavioral analyses

Mean absolute recall error was calculated for each subject and each condition. A two-tailed paired t-test was conducted to test whether performance differed across conditions.

We applied a variant of the target confusability competition (TCC) model to track the fate of the IMI (Schurgin et al., 2020). With TCC models, the subject’s report on a trial is assumed to be based on an aggregated familiarity landscape that reflects the influence of all of the items currently in WM (see Fig. 2A for an example illustration). TCC estimates the memory strength of an item with a single parameter, d’, which represents the magnitude of the signal corresponding to that item. Whereas the original formulation of the TCC model was restricted to the items currently in WM, and d’ was bounded to be non-negative, for the present study we extended this model to include an estimate of d’ for the putatively removed item (c.f., (Zhang and Lewis-Peacock, 2023)), and so we needed to modify it in order to allow d’ to take on negative values. In particular, for the ABC-retrocuing task, although the passive removal of the IMI was predicted to leave a small but positive d’ (corresponding to an incompletely removed memory trace), the active removal of the IMI was predicted to produce a negative d’. This is because, according to the hijacked adaptation model, the active removal of the IMI is accomplished by down-modulating the gain of sensory channels tuned to its orientation. Thus, the subsequent encoding of item *C* should be influenced such that the final familiarity landscape would be reduced for orientations close to the IMI (see Fig. 2B). A negative IMI d’ in the overlap condition would be consistent with this model.

**Figure 2.**
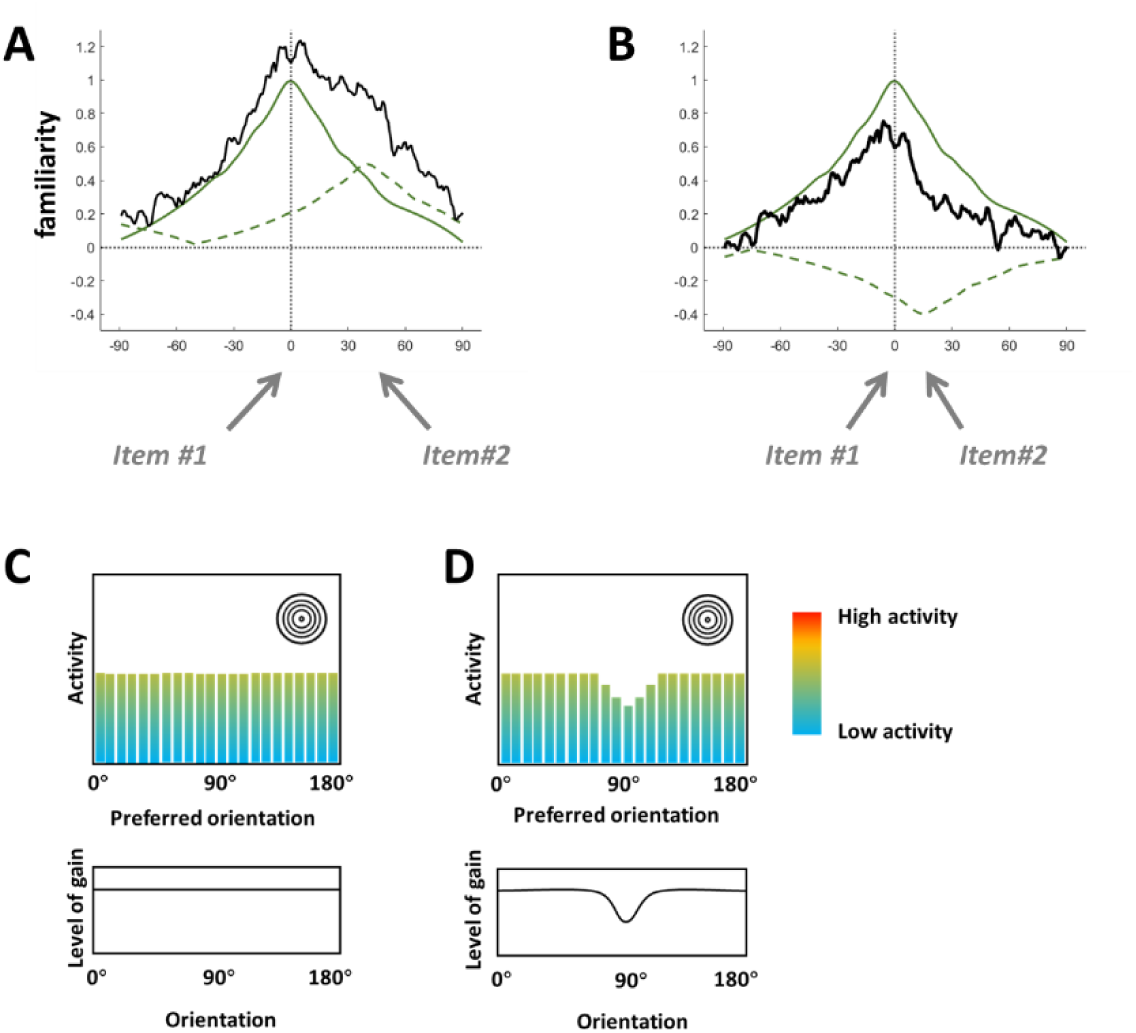
Illustrations of predictions of the hijacked adaptation model. A. Hypothetical behavioral data from a single trial, fit to TCC model, in which the familiarity signals for item #1 (solid green line) and for item #2 (dashed green line, are strong and weak, respectively. The solid black curve is the resultant familiarity landscape that is accessible to the cognitive system, generated by taking the sum of two green curves and adding some noise (note the “shoulder” corresponding to the weaker item #2). B. Hypothetical behavioral data from a trial in which item #2 has been actively removed, the down-modulation of its gain resulting in negative familiarity centered on its orientation (note the consequent reduction in the corresponding portion of the familiarity landscape). C. On trials in which neither item has been actively removed, the gain of all orientation-tuned sensory channels is comparable (lower plot), and so a nonspecific visual “ping” has the effect of activating each by the same amount (upper plot; each bar illustrates the level of activity of an orientation-tuned sensory channel). D. On trials in which an item (90°) has been actively removed, the gain of sensory channels centered on that item’s orientation has been reduced. Consequently, the response of these sensory channels to a nonspecific visual “ping” is lower.

The TCC model incorporates the psychological similarity between stimulus items, and posits that a graded familiarity signal is generated by each item in WM (Schurgin et al., 2020). For orientation, a 0° memorandum would boost familiarity for all possible orientations ranging from -90° to 90°, with the magnitude of this boost for any one value determined by its similarity to 0° (e.g., Fig. 2A). To apply the TCC model, we used the psychometric similarity function for orientation estimated in a previous study (Cai et al., 2022). To assess whether active removal of the IMI influenced the processing of item *C*, we focused on the 50% of trials in which subjects were probed to recall item *C*. The familiarity landscape for each trial was modeled to be the sum of the familiarity of the probed item (i.e., item *C*, estimated with probed d’) and the signal corresponding to the IMI (estimated with IMI d’). Both probed d’ and IMI d’ were allowed to take a negative value. To calculate d’ values, we combined the data from all subjects into a “super subject” and conducted all TCC analyses on this super subject. The TCC model was fitted to data from the overlap condition and the no-overlap condition separately, using Markov Chain Monte Carlo (MCMC function in MemToolbox; (Suchow et al., 2013)). 15,000 post-convergence samples were taken to calculate the 95% confidence interval (95% CI) of the parameter estimates. To investigate the difference between conditions, the differences in the estimated parameters from the 15,000 samples in each condition were calculated and the 95% CI was generated from them. *p* values were also calculated from each pool of samples by calculating the proportion of samples higher or lower than 0 and multiplying the result by 2 to make it two-tailed.

### EEG recording and preprocessing

Scalp EEG was recorded from 60 electrodes with an actiCHamp Plus system (Brain Products), at a sampling rate of 1,000 Hz. The position of electrodes was based on the extended 10-20 international system. The recording was referenced online to a frontal electrode (FCz). Preprocessing and analysis were conducted with EEGLAB (Delorme and Makeig, 2004), Fieldtrip (Oostenveld et al., 2011) and custom MATLAB scripts. Data were first down-sampled to 500 Hz and then a bandpass filter of 1 – 100 Hz was applied. Bad channels were detected and removed using the pop_rejchan function of EEGLAB. Then, the data were re-referenced to the average of all remaining electrodes. Continuous EEG data were segmented into 13.25-sec epochs (from 1.25 sec before sample onset to the offset of the feedback) and epochs containing artifacts were detected and rejected using the pop_autorej function of EEGLAB. After reducing the dimensionality of data to 32 with principal component analysis, and applying an independent component analysis, components containing eye-movement artifacts, muscle artifacts and line noise were detected and removed using ICLabel (Pion-Tonachini et al., 2019). Finally, bad channels were interpolated.

### Logic of EEG analyses

As a mechanism of top-down control, one would expect hijacked adaptation to have two discrete sets of neural correlates: one associated with the triggering of the control signal, tightly timelocked to the prioritization cue, and one associated with the consequences of the deployment of this control signal, which would be revealed in ping-evoked activity. Thus, we focused on activities timelocked to these two events for our EEG analyses.

In activity timelocked to the onset of the retrocue, we planned to look for evidence of the top-down control signal that is hypothesized to implement hijacked adaptation. We hypothesized that one potential manifestation of this signal might be traveling waves, the spatial propagation of neural oscillations across the cortex that have been shown to support communication between brain regions, and that have been proposed to subserve cognitive processing (Muller et al., 2018; Xu et al., 2023; Luo and Ester, 2024). In particular, forward (posterior-to-anterior) and backward (anterior-to-posterior) traveling waves have been shown to play an important role in bottom-up and top-down processing of information, respectively (Alamia and VanRullen, 2019; Mohan et al., 2024). Thus, analyses time-locked to the retrocue were predicted to reveal a stronger backward traveling wave on overlap versus on no-overlap trials.

Regarding the consequence of active removal, the down-regulated gain of sensory channels tuned to the to-be-removed orientation was predicted to reduce the overall excitability of the perceptual circuit responsible for its processing (see Fig. 2D for an illustration). To assess this predicted effect, we delivered a strong orientation-nonspecific visual ping during the delay period that followed the retrocue, so as to trigger a robust evoked response and thereby detect the change in excitability of the critical perceptual circuit. Specifically, we predicted that the ping-evoked response would be weaker following active removal of the IMI (Fig. 2D), relative to passive removal (Fig. 2C), because only the former would be accompanied by a gain reduction in orientation-tuned sensory channels corresponding to the actively removed item. For traveling waves timelocked to the ping, we expected to see decreased forward traveling waves on overlap versus on no-overlap trials, corresponding to the decreased bottom-up processing of information with hijacked adaptation.

### ERP analyses

We studied the ERPs evoked by the onset of the retrocue and the ERPs evoked by the ping. For all ERP analyses the EEG was baseline-corrected using the 750 ms before the start of the trial. To test the difference of retrocue-evoked ERPs across two conditions, we applied the cluster-based permutation test in Fieldtrip. First, a paired t-test was conducted and all data points with a p-value lower than 0.05 were identified. Temporally and spatially adjacent significant data points were taken into the same cluster and the summed t value for each cluster was calculated. Finally, the summed t values were compared to the largest summed t values generated from 1,000 permutated data with randomly permuted labels of conditions, and significant clusters were detected with a two-tailed alpha of 0.05.

For the ping-evoked response, we focused on posterior sensors (P7, P8, P5, P6, PO7, PO8, PO3, PO4, POz, O1, O2, Oz), because the hijacked adaptation model predicts a change in the sensory circuits responsible for encoding stimulus orientation. To analyze the statistical significance, temporal-cluster-based permutation tests were conducted with 1,000 random permutations and a two-tailed alpha of 0.05.

### Traveling wave analyses

Forward (posterior-to-anterior) and backward (anterior-to-posterior) traveling waves were assessed in EEG epochs timelocked to retrocue onset and timelocked to ping onset. Traveling waves were estimated with a procedure based on 2D fast Fourier transform (2D-FFT). We used four anterior-posterior axes linking frontal and occipital electrodes (Fig. 5A; [PO7, P5, CP5, C5, FC5, F5, AF3], [O1, PO3, P3, CP3, C3, FC3, F3], [O2, PO4, P4, CP4, C4, FC4, F4], and [PO8, P6, CP6, C6, FC6, F6, AF4]). For each axis of electrodes, a 500 ms time window was used and the voltage from electrodes was taken to construct a 7 (channels)-by-250 (timepoints) image. Then, a 2D-FFT was applied to this image. The upper and lower quadrants of the resulting spectra quantify the magnitude of the backward (anterior-to-posterior) and forward (posterior-to-anterior) traveling waves, respectively (Alamia et al., 2023; Luo and Ester, 2024), with the x axis corresponding to the temporal frequency of the traveling waves and the y axis corresponding to their propagating speed. Then, for each temporal frequency, the maximum value in each quadrant was extracted. This resulted in a vector of traveling wave power for this time window.

Then, the time window was slid by 50 ms and the procedure repeated. As the result, a time-by-frequency matrix of traveling wave power was generated for each axis of electrodes and each direction (forward versus backward).

The power matrix was corrected with a baseline matrix. The baseline matrix was constructed by randomly shuffling electrodes and generating time-by-frequency matrices with the procedure described above, repeating this process 50 times, then averaging the results. Lastly, traveling wave power was calculated as a decibel ratio between the observed and baseline power:

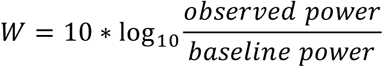

Forward and backward waves were estimated separately from epochs timelocked to retrocue onset and timelocked to ping onset. (Note that these analyses allow for forward and backward waves to be present along the same axis during the same temporal epoch.) For the comparison between conditions, traveling wave power from the four axes of electrodes was averaged and cluster-based permutation testing (with 1,000 random permutations) was used to detect significant clusters.

### Statistical Analyses

Details about the statistical test used for each analysis appear in preceding subsections of the Methods. In brief, a two-tailed paired t-test was used to test whether there was a difference in behavioral performance across conditions; for the TCC model analysis, the 95% CI of the parameter estimates was calculated based on the Markov Chain Monte Carlo post-convergence samples and the statistical significance of difference between conditions was also obtained from these samples; for ERPs analyses and traveling wave analyses, clustered based permutation tests were run to detect significant differences between conditions.

## Results

### Behavioral Results

Mean absolute recall error indicated compliance with task instructions (SD = 12.540±3.069 deg), with recall error in the overlap condition (SD =11.960±3.117 deg) significantly lower than in the no-overlap condition (SD=13.120±3.195 deg; t(24) = 3.8452, *p* < 0.001).

Applying the TCC model to trials in which subjects were probed to recall item *C* indicated that this item evoked a strong familiarity signal in both conditions (Fig. 3; no-overlap probed d’ = 2.532, 95% CI = [2.462, 2.604], *p* < 0.0001; overlap probed d’ = 2.864, 95% CI = [2.785, 2.946], *p* < 0.0001). The probed d’ was significantly higher in the overlap condition than in the no-overlap condition (Difference CI = [0.226, 0.448], p < 0.0001). In the no-overlap condition the IMI d’ was small but significantly higher than zero (Fig. 3; no-overlap IMI d’ = 0.252, 95% CI = [0.159, 0.341], *p* < 0.0001), suggesting that a residual trace of IMI persisted at the end of the trial. In the overlap condition, IMI d’ was numerically negative (overlap IMI d’ = -0.096, 95% CI = [-0.208, 0.016], *p* = 0.0888), and significantly lower than the no-overlap IMI d’ (difference CI = [-0.4925, -0.2046], *p* < 0.0001).

**Figure 3.**
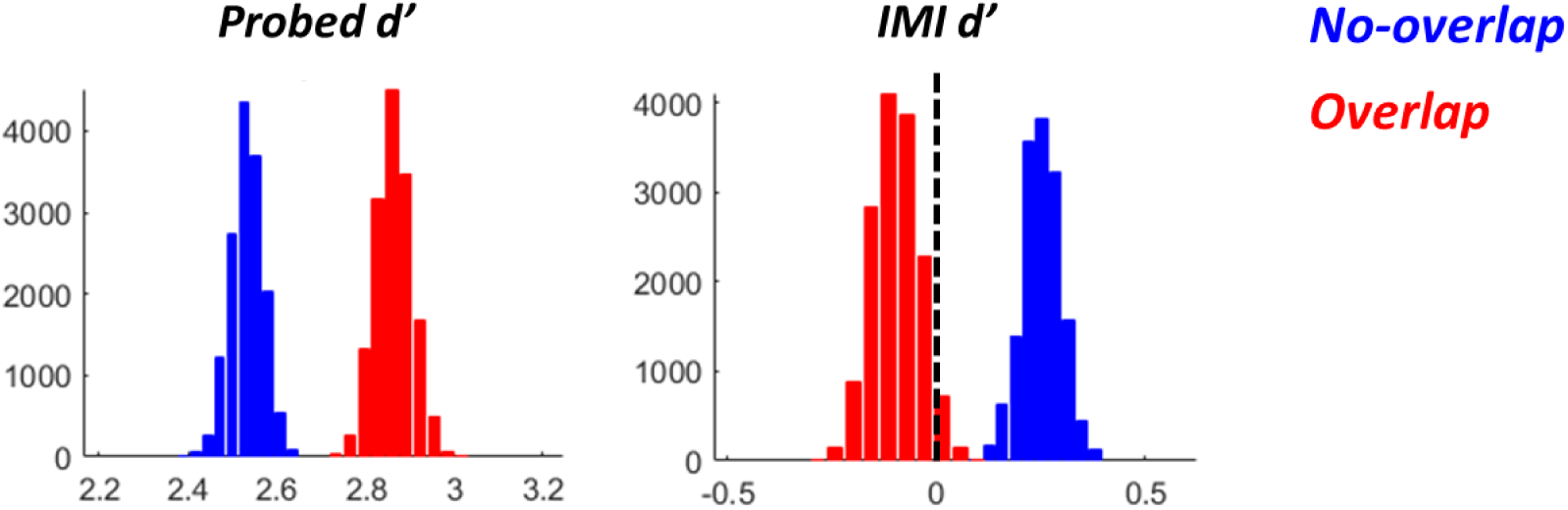
Histogram of parameter estimates from the 15,000 post-convergence samples with TCC model, of trials on which item C was probed. Note the difference, for item C (“probed d’”) vs. for the IMI (“IMI d’”), in values on the horizontal axis.

### EEG results

#### Retrocue-evoked ERPs

In both conditions, scalp topographies showed a large negativity at central midline electrodes. Comparison of ERPs averaged over these 8 central midline electrodes (Fz, F1, F2, FC1, FC2, C1, Cz, C2) revealed that the magnitude of the ERP was larger in the overlap condition, the two conditions diverging beginning during the initial negative-going deflection following retrocue onset and this difference persisting for roughly 250 msec (Fig. 4; 208 to 456 ms after retrocue onset, *p* = 0.002).

**Figure 4.**
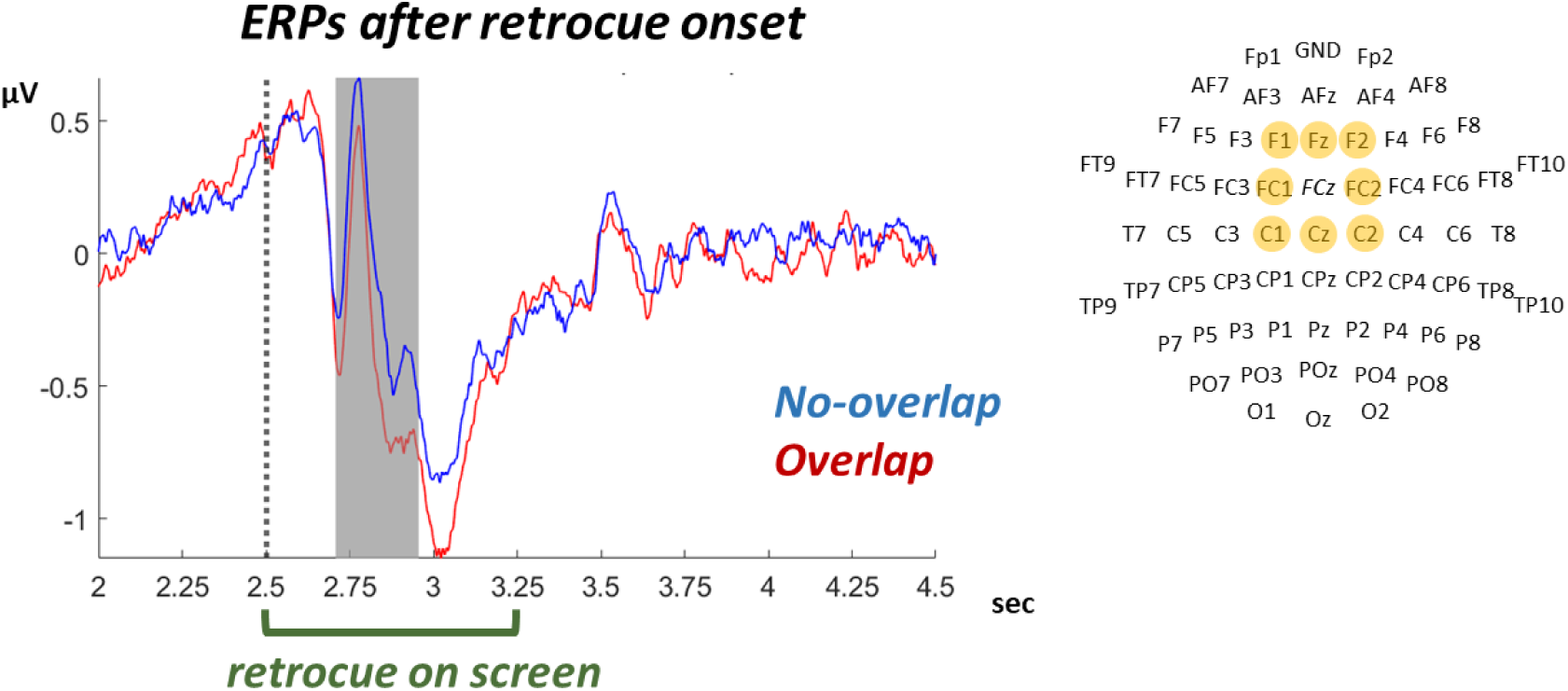
ERPs from frontal midline electrodes (see right panel), timelocked to retrocue onset. Gray shaded area indicates time with significant difference across conditions.

#### Traveling waves timelocked to the retrocue

The forward traveling wave analyses revealed evidence for tonically elevated forward wave at low frequencies (< 10Hz), and a tonically suppressed forward wave in the beta band (20-26 Hz), in each condition, and for each spanning the duration of Delay 1, the retrocue, and Delay 2.1. In both conditions there was a brief increase in the magnitude of the low-frequency forward wave that was centered on the offset of the retrocue (i.e., at time 3.25 sec). The two conditions only differed for a brief period starting from ∼800 ms after retrocue onset, when the tonically suppressed forward traveling in the beta-band dipped lower in the no-overlap condition relative to the overlap condition (Fig. 5D left, *p* = 0.01).

**Figure 5.**
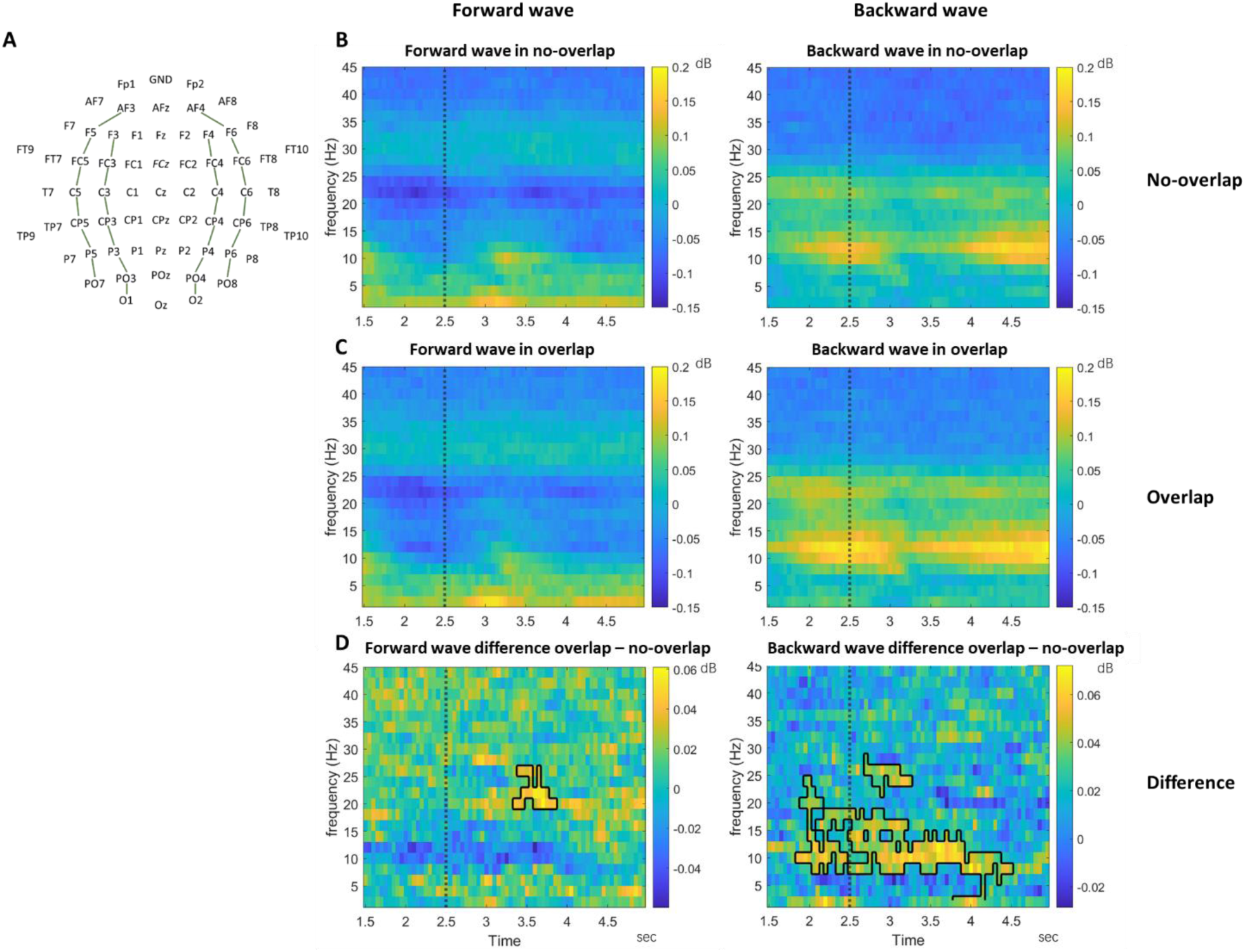
Forward and backward traveling waves in EEG, assessed from the average over four axes of electrodes (A), and timelocked to the retrocue onset (dashed line at 2.5 sec), in no-overlap (B) and overlap conditions (C). D shows the difference between conditions with significant clusters marked with black contours. The left column is forward waves and the right column shows backward waves.

Backward traveling-wave analyses revealed similar patterns in both conditions of persistently elevated magnitudes in two frequency bands: one spanning high-alpha/low-beta and one, less strong, at higher frequencies in the beta band, spanning from ∼20-25 Hz (Fig. 5B right and 5C right). (Note that we cannot rule out the possibility that the higher-frequency of these two backward traveling waves may be a harmonic artifact of the lower-frequency one.) In both conditions these backward traveling waves were prominent prior to retrocue onset. The power of these backward waves was stronger in the overlap condition both during the epoch surrounding the retrocue (in alpha/low-beta from ∼700 ms before retrocue onset until ∼2 sec after retrocue onset, Fig. 5D right, *p* = 0.002) and in higher beta beginning shortly after the retrocue onset (∼200-750 ms after retrocue onset, Fig. 5D right, *p* = 0.03)

#### Ping-evoked ERPs

For both conditions the ping evoked a large response in posterior electrodes, with a larger negative-going amplitude for the no-overlap condition beginning with the first negative-going deflection and persisting for roughly 135 msec (Fig. 6; 220 to 356 ms after ping onset, *p* = 0.028). To assess whether this difference is best characterized as a larger-amplitude ping-evoked response or a DC shift between the two conditions, we calculated, for each subject, the peak-to-peak distance between the initial negative-going deflection and the subsequent large-amplitude positive-going deflection, reasoning that if the difference were due entirely to a DC shift, the peak-to-peak distances for the two conditions would be the same. For this analysis, the negative peak was defined as the lowest voltage in a 200-msec time window centered on the group-average negative peak and the positive peak was defined as the highest voltage in a 200-msec time window centered on the group-average positive peak.

**Figure 6.**
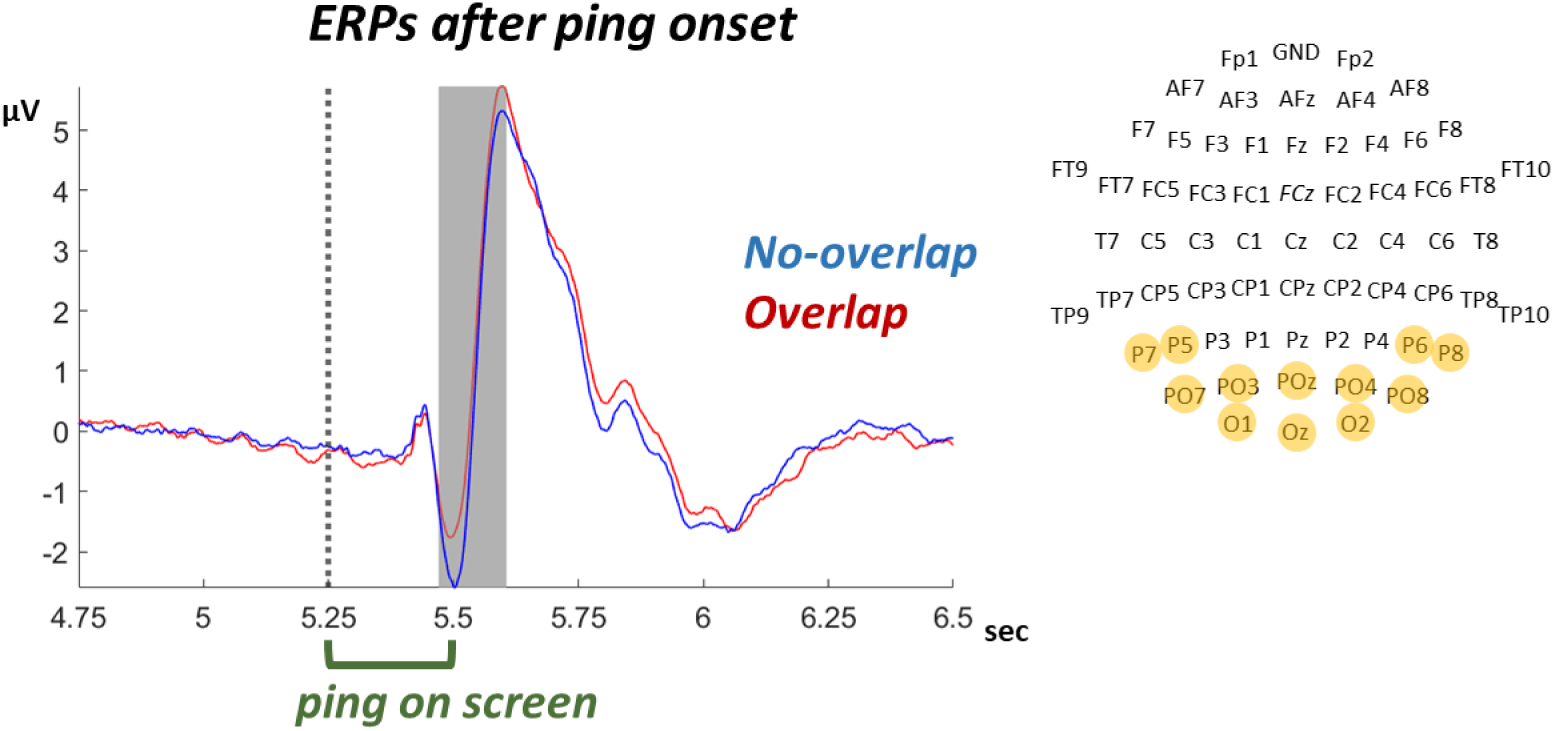
Ping-evoked ERPs from posterior electrodes (see right panel). Gray shaded area indicates time with significant difference across conditions.

Statistical comparison of peak-to-peak distance between conditions indicated that the difference approached, but did not achieve, the threshold for significance (t(24) = -1.9789, *p* = 0.0594). Although this result was equivocal, it approached the level at which one would say that this difference was due, at least in part, to a larger-amplitude ping-evoked response.

#### Traveling waves timelocked to the ping

Forward traveling waves, when timelocked to the onset of a visual stimulus, are believed to correspond to bottom-up processing of the perceptual input (Alamia and VanRullen, 2019). Consequently, the hijacked adaptation model predicted that the forward traveling wave triggered by the ping would be reduced during blocks that encouraged active removal. Consistent with this prediction, the visual ping evoked a strong forward traveling wave in the theta band (∼4-8 Hz), starting from ∼250 ms after ping onset in both conditions (Fig. 7A left and 7B left). This traveling wave was numerically weaker in the overlap condition relative to the no-overlap condition (Fig. 7C left).

**Figure 7.**
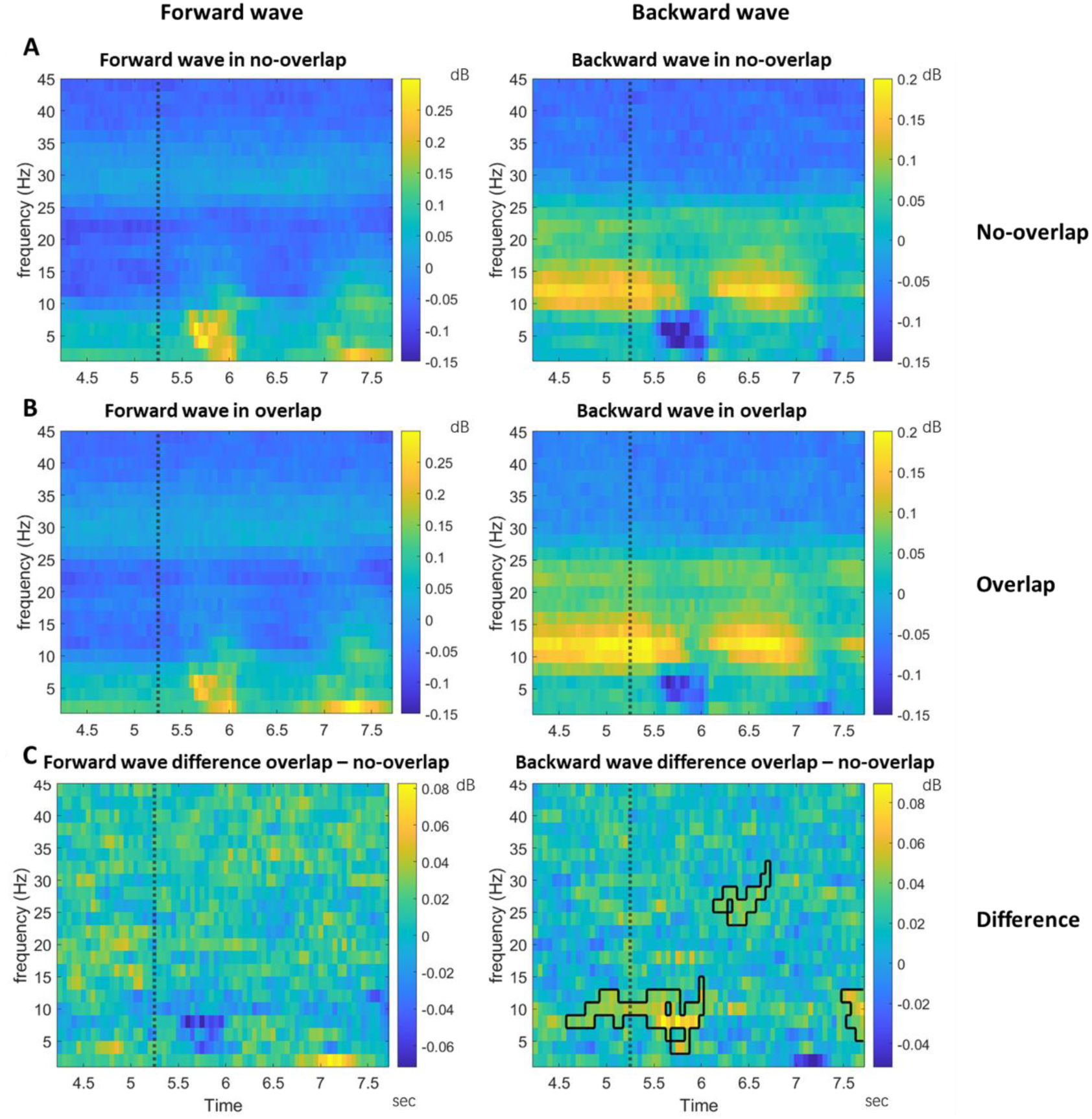
Forward and backward traveling waves in EEG timelocked to the ping onset in the no-overlap (A) and overlap conditions (B). Row C shows the difference between conditions. The left column shows forward waves and the right column backward waves. Note that the scale is different for forward and backward traveling waves. The significant difference clusters around or after 6.5 sec in time were likely caused by the onset of item C, which was at 6.5 sec. Plotting conventions are the same as figure 5.

A tonically elevated backward traveling wave spanning high-alpha/low-beta frequencies (∼10-15 Hz) and extending in time from Delay 2.1 until well into Delay 3, with a pause briefly following the ping, was prominent in both conditions (Fig. 7A and B). This was likely a continuation of the backward traveling wave observed in the analyses timelocked to the retrocue (Fig. 5B anc C). The power of this backward waves was stronger in the overlap condition during the epoch surrounding the ping (in alpha/low-beta starting from ∼650 ms before ping onset until ∼750 ms after ping onset, Fig. 7C right, *p* = 0.002). A significant difference was also observed in high beta (∼23-28 Hz) during the time corresponding to the onset of item C, which was at 6.5 sec Fig. 7C right, *p* = 0.016).

## Discussion

Although it is widely assumed that updating the contents of working memory can include the operation of actively removing no-long-relevant contents (e.g., (Jonides et al., 1997; Postle et al., 2001)), the mechanisms that might accomplish this operation have received far less attention than other aspects of working memory. We approached this question with the assumption that there may be (at least) two qualitatively different ways whereby information leaves working memory: passively, as a consequence of the withdrawal of attention (Chatham and Badre, 2013; Barbosa et al., 2020), or actively, via the application of cognitive control. (For alternative accounts, see (Kim et al., 2020; Beukers et al., 2021).) We operationalized these two scenarios with a single task in which we manipulated the level of cue conflict between a no-longer-relevant item and a newly added item, reasoning that the low-conflict (“no-overlap”) condition would encourage passive removal, whereas the high-conflict (“overlap”) condition would encourage active removal. In previous work, using this, and similar, designs, we have generated indirect evidence that the no-longer-relevant item is, indeed, processed differently in the two conditions. More specifically, the fact that, in the high-conflict condition, this item exerts a repulsive serial bias is consistent with the idea that active removal might be accomplished via a mechanism of hijacked adaption (Shan and Postle, 2022; Teng et al., 2022). The hijacked adaption model posits that the removal of information from WM is accomplished by a top-down control signal that down-modulates the gain of perceptual circuits tuned to that item. In the present study we have extended the previous work by finding direct behavioral evidence for the suppression of the target item in the active-removal condition, and EEG effects consistent with the implementation of hijacked adaptation (at the time of the retrocue) and with a consequence of hijacked adaption (later in the trial, at the time of the ping).

The hijacked adaptation model posits that the active removal of an item in working memory (the irrelevant memory item; IMI) is accomplished by the down-modulation of gain in perceptual circuits tuned to that item. This particular mechanism has been posited, rather than some other form of inhibition, to account for the residual effects of active removal on the processing of subsequent items.

For example, whereas the IMI exerts an attractive bias on recall on the subsequent trial when it has been passively removed, the sign of this serial dependence effect is reversed after active removal (Shan and Postle, 2022; Teng et al., 2022). Here we provide novel behavioral evidence for this account with a modified TCC analysis that allowed us to estimate the effect of active removal of the IMI during that same trial. TCC models have been previously used to study the memory strength of the memorandum (Schurgin et al., 2020) and the intrusion effect of a distractor (Zhang and Lewis-Peacock, 2024). Here we further extended the model in order to study the influence of the IMI, by including the IMI in the model and by allowing d’ to take on negative values. This allowed us to make the novel observation that active removal of the IMI leads to a drop in the familiarity landscape centered on the orientation of the IMI, with d’ for the IMI taking on a numerically negative value. In the no-overlap condition, in contrast, the lMI left a bump in the familiarity landscape, indicating it was not fully removed from WM, and the positive value of d’ was significantly different from its value in the overlap condition. This pattern in the no-overlap condition was in line with our expectation for passive removal, and consistent with many previous findings (Monsell, 1978; Fischer and Whitney, 2014; Bae and Luck, 2019; Samaha et al., 2019). Thus, these results represent novel behavioral evidence for active versus passive removal from WM.

At the neural level, our design incorporated a visual ping to detect the predicted difference in perceptual circuits after subjects conducted active versus passive removal. Because the hijacked adaptation model predicts decreased gain in IMI-tuned perceptual channels, the overall response to the ping was expected to be reduced on overlap versus no overlap trials. Two analyses of the ping-evoked EEG activity supported this prediction. First, the ERP evoked by the ping was significantly weaker in the overlap condition compared to the no-overlap condition, with the divergence in ERPs beginning ∼250 ms after ping onset, suggesting a reduction of perceptual-circuit excitability after subjects conducted active removal. Second, a traveling-wave analysis revealed a numerical reduction of the forward traveling wave, in the theta band, also starting at ∼250 ms after ping onset. Forward traveling waves have been proposed to index the feedforward processing of visual inputs (Alamia and VanRullen, 2019; Mohan et al., 2024), possibly as a manifestation of the synchronization of neural oscillations between brain regions (Fries, 2015)). Together, these findings provide novel evidence that a mechanism for the active removal of information from WM, presumed to be initiated once the retrocue designated the IMI, had the effect of suppressing the visual processing of a task-irrelevant ping presented 2 sec later in the trial.

We acknowledge that a more direct evaluation of the hypothesized mechanism of hijacked adaptation would be via measurements of the neural representations of the information being removed from WM, such as via multivariate reconstruction methods like inverted encoding modeling (IEM; c.f., (Lorenc et al., 2020; Yu et al., 2020)). However, our attempts at reconstruction of stimulus orientation were unsuccessful, even for items actively maintained in WM (data not shown). This was likely due to the fact that our stimuli were presented at locations in the periphery, and because there were always two items in WM. (For a direct illustration of the effects on stimulus representation of hijacked adaptation, see (Teng et al., 2023))

Active removal via hijacked adaptation is assumed to be accomplished via top-down signaling that commands the down-modulation of gain in posterior perceptual circuits. In the ABC-retrocuing task, this control signal is assumed to be triggered by the retrocue, which designates (by implication) that trial’s IMI. As with the ping, we carried out two sets of analyses timelocked to the onset of the retrocue: ERP, and traveling wave. For ERPs, we found that subjects showed a stronger negative-going deflection of the ERP at frontal-midline electrodes for the overlap condition relative to the no-overlap condition. For traveling waves, we observed widespread and prominent backward waves that were stronger in the overlap than the no-overlap condition. These backward traveling waves were strongest in a frequency band spanning alpha and low beta, started prior to retrocue onset, and persisted until 2 sec after the retrocue onset. Thus, the backward waves found in the current study could be reflecting a top-down signal that exerts cognitive control over posterior brain areas (c.f., (Alamia and VanRullen, 2019; Alamia et al., 2023; Luo and Ester, 2024)). At a general level, this aligns well with literatures showing the important role of alpha- and beta-band synchronization in feedback signaling (Bastos et al., 2015; Fries, 2015; Das and Menon, 2021, 2022). Although we hesitate to speculate too much about the specific function(s) of the pattern of backward traveling waves found in the current study, it may be that the stronger alpha/low-beta-band backward traveling waves in the overlap condition correspond to a preparatory control process that starts before retrocue onset and persists until removal of the IMI is completed. In a set of WM tasks, the magnitude of alpha backward traveling waves has been found to be modulated by the load of WM such that higher WM load was associated with weaker alpha backward waves (Zeng et al., 2024). The authors proposed that alpha backward waves may reflect top-down inhibitory gain control (Zeng et al., 2024), which aligns well with our results in the current study.

Oscillations in the beta band, on the other hand, are believed to be important for implementing domain-general inhibitory control (Wessel and Anderson, 2024), including for the inhibition of WM content (Lundqvist et al., 2024). Although our analyses also detected a significant difference, across conditions, in forward waves in retrocue-locked activity, it is unclear how to interpret this, because this effect was driven by greater suppression of forward traveling waves in the no-overlap condition (i.e., lower amplitude in comparison to baseline, see Methods).

To summarize, by using EEG recordings and TCC modeling of behavioral data, we have shown that active removal of information from WM is associated with a top-down control process manifesting at frontocentral electrodes. It results in reduced excitability at posterior electrodes (a hypothesized correlate of down-modulation of gain in posterior perceptual circuits) and behavioral evidence for a stimulus-specific reduction in the familiarity landscape from which the recognition decision is read out. Together, they add to growing evidence for hijacked adaptation as a mechanism for the active removal of information from WM.

## Acknowledgments

This research was funded by National Institutes of Health Grant MH131678 (to B.R.P.).

